# Living fast, dying young: anthropogenic habitat modification influences the fitness and life history traits of a cooperative breeder

**DOI:** 10.1101/2023.05.13.540639

**Authors:** Alejandro Alaman, Enrique Casas, Manuel Arbelo, Oded Keynan, Lee Koren

## Abstract

Modification of natural habitats can lead to an ecological trap, as animals can maladaptively select lower-quality habitats over natural landscapes. Modified habitats reduce group size and prevalence, the success of dispersing individuals, and breeding success. However, little is known about how these effects on fitness affect the sociality of cooperative breeding species, which may be particularly susceptible to habitat modification. Here we show that the selection of anthropogenically modified habitats negatively affects the fitness, which has consequences in the life history traits of a cooperative breeder.

We used data collected during six years from a monitored population of Arabian babblers (*Argya squamiceps*) and found that groups living in modified habitats breed more than those in natural habitats but that juvenile and adult survival in modified habitats was lower. Individuals living in modified habitats acquired dominance earlier than individuals from natural habitats. Males in modified habitats also dispersed earlier than those from natural habitats.

Our results suggest that modified habitats entail an ecological trap for Arabian babblers, which adjusts their life history traits as a response. Understanding the adaptation of social and cooperative breeding species to habitat modification is the first step for mitigating the processes behind human-related population declines.

## Introduction

Humanity’s impact on the earth’s ecosystems in the last tens of thousands of years has been so extensive that it is now defined as a geological era - the Anthropocene^1^. The last global assessment report on biodiversity and ecosystem services^2^ estimated that about 75% of the earth’s land had been modified by humans and that more than a third had been converted to crop or livestock production, which is the main cause of the rapid decrease in biological diversity and the intense extinction of wild populations^3,4^. The impact of habitat modification on biodiversity loss and population extinction has been mostly studied in temperate and tropical ecosystems^5–8^. However, the consequences of habitat loss and modification in arid ecosystems may be more profound^9^.

Arid environments are generally resource-poor habitats, so agricultural activities that add water and fertilizers are expected to radically change it. New resources attract native species^10–12^ at the cost of reduced survival due to increased roadkill^13^, higher predation^14,15^, and chemical exposure^16,17^. Thus, intensive agriculture in arid environments constitutes an ecological trap, as the cues that animals use to assess habitat quality and the true quality of the area are decoupled^18–20^. The more attractive the new environment is for individuals, and the more reason for it to potentially reduce survival or reproduction, the stronger the ecological trap^19^.

Cooperative breeders, whereby individuals delay their dispersal and help care for offspring that are not their own^21^, are mostly found in areas of environmental uncertainty^22^. Since environmental conditions are known to affect the costs and benefits of cooperation^23–27^, they may be especially vulnerable to anthropogenic changes and ecological traps. The ecological constraint hypothesis^28–30^ suggests that the evolutionary causes of delayed dispersal in cooperative breeders are due to a lack of suitable territories, low survival rates during dispersal, low probability of mating, and low reproductive success^24,28,29^. Habitat modification have negative demographic effects in cooperatively breeding species, including reduced group size, fewer dispersers, lower breeding success, and higher mortality^31–34^. Group density and/or population size declines can lead to a demographic Allee effect^35–37^, i.e., a positive effect of population size or density on group performance or population growth, so that small populations or populations with a low group density are more likely to become extinct^35^.

Arabian babblers (*Argya squamiceps*) are group-living, territorial, obligate cooperatively breeding birds found along the Arabian desert and the Sinai Peninsula^38^. Juveniles delay dispersal for an average of 1–3 years (but sometimes completely) whilst helping their parents care for their siblings. Reproductive skew is high; each group usually contains only one breeding pair (> 96% of groups). Mortality rates are the highest during the first year of life^39^. Adults remain in their natal group as helpers until they acquire dominance, disperse, or die. Dispersal is female-biased and can be alone or in a coalition of several individuals^39,40^, in order to join or establish a new group or become a floater (individuals with no fixed territory, usually found alone or in small coalitions^41^). Over the > 50 years of Arabian babbler study by Amotz Zahavi et al. in our study site, the agricultural land has expanded, and groups of Arabian babblers began to inhabit it. We observed that groups living in natural and modified habitats showed different preferences in the selection of nesting and roosting trees. Groups in natural areas use mostly native trees such as Umbrella thorn acacia (*Vachellia tortilis*) or Christ’s thorn jujube (*Ziziphus spina-christi*), while groups in modified habitats combine these species with others of anthropogenic origin, as mango (*Mangifera indica*) and date (*Phoneix dactilifera*). Plantations are also used by these groups for foraging or as a water source. This increase in resources could be expected to benefit groups which inhabit this habitat, however, our preliminary exploration of the data did not support this hypothesis. Therefore, we suspect that the agriculture-modified habitat could entail an ecological trap, attracting individuals and groups to it, yet reducing their fitness. Under this scenario, the aim of this study is to explore the effect of habitat choices of Arabian babbler on their fitness and the consequences that might have on their life history traits and sociality.

## Results

We used six years of data (2016-2021) of 572 individuals from 21 different groups in our monitored population of Arabian babblers to evaluate whether life history traits and demography are different between natural habitats and anthropogenically modified habitats. Allee effect found in Arabian babblers suggests that habitat degradation and human disturbance are main disturbances^37^. During this period, we followed a total of 203 breeding events from the study groups. We monitored breeding success, juvenile and adult survival rates, and their effect on group size. We also compared two life history traits: age at dominance acquisition and age at dispersal. We used information about the presence of individuals, monitored nests and identified and characterized habitats using Remote Sensing (RS) products and images to evaluate anthropogenic impacts on fitness and life history traits (Fig. 1).

**Figure 1.**
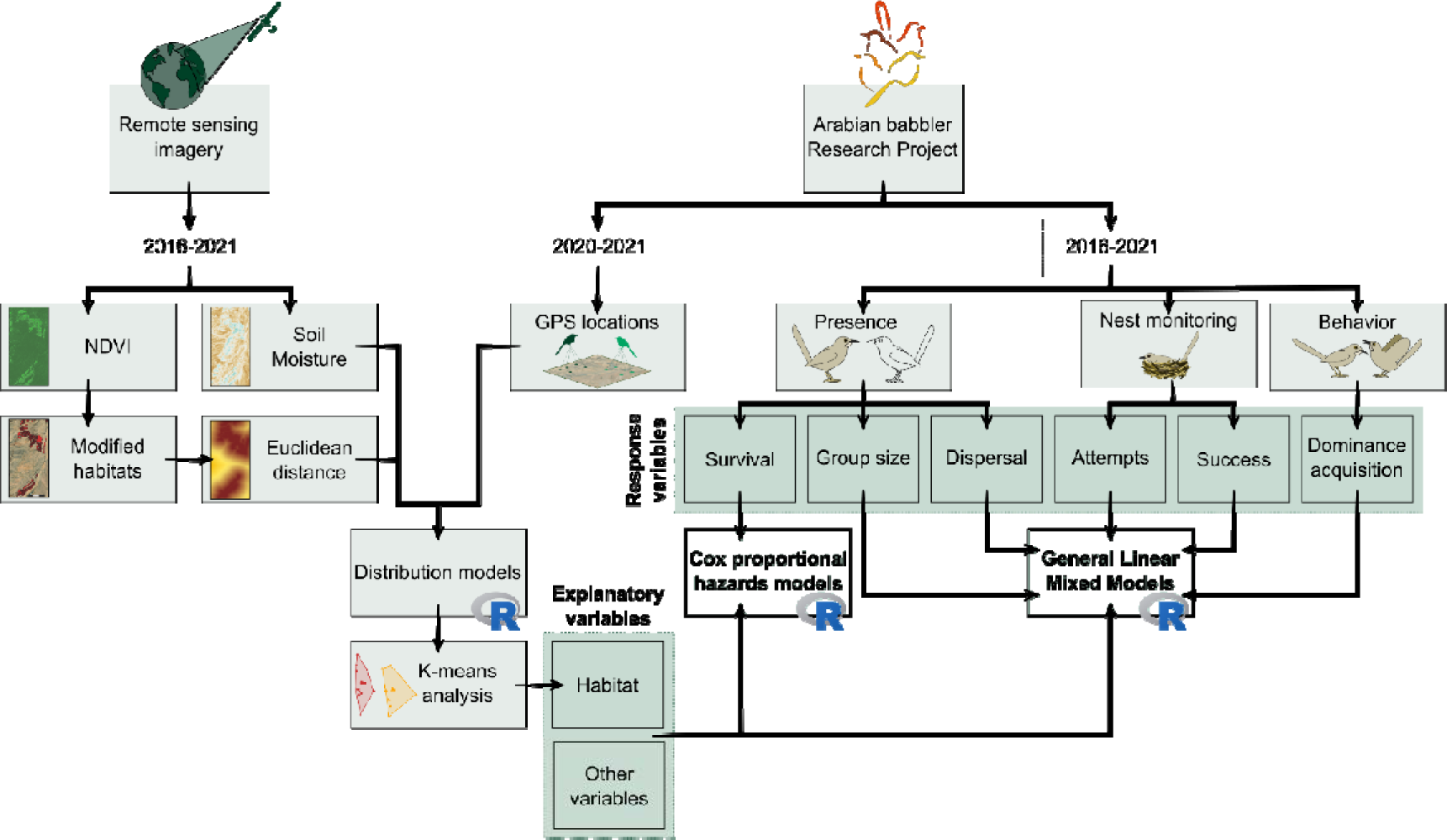
Overview of the study design. The data for this study are from the Arabian babbler Research Project, which has been ongoing since 1971, and from available remote sensing imagery extracted from Copernicus Sentinel 1 (S1) and Sentinel 2 (S2). The oldest imagery data available for the region was from the 2016 breeding season, which was set as the beginning of this study. From the Arabian babbler Research Project we had individual GPS locations (collected since 2020), presence (from which we calculated survival, group size, and dispersal), nest monitoring (from which we calculated attempts and success) and behavior (from which we observed dominance acquisition). From the remote sensing imagery we obtained data for soil moisture (S1) and NDVI (S2). NDVI was used to characterize modified habitats areas and Euclidean distance to them. With Sild moisture, Euclidean distance to modified habitats and group locations we created distribution models for territory characterization of each group. We used K-means analysis to classify the groups in modified and natural habitats. We used habitat and other variables to analyze the effect of habitat on survival using Cox proportional hazards models. The effect of habitat on group size, dispersal, breeding attempts and success, and dominance acquisition was analyzed via General linear mixed models.

### Habitat characterization

#### Species distribution models (Group spatial distribution)

Generalized Additive Models (GAM) presented strong predictive capabilities with an Area Under the Curve (AUC) value of 0.946, sensitivity value of 0.93, and 0.916 for specificity (e.g., projected group territories Fig. 2A).

**Figure 2.**
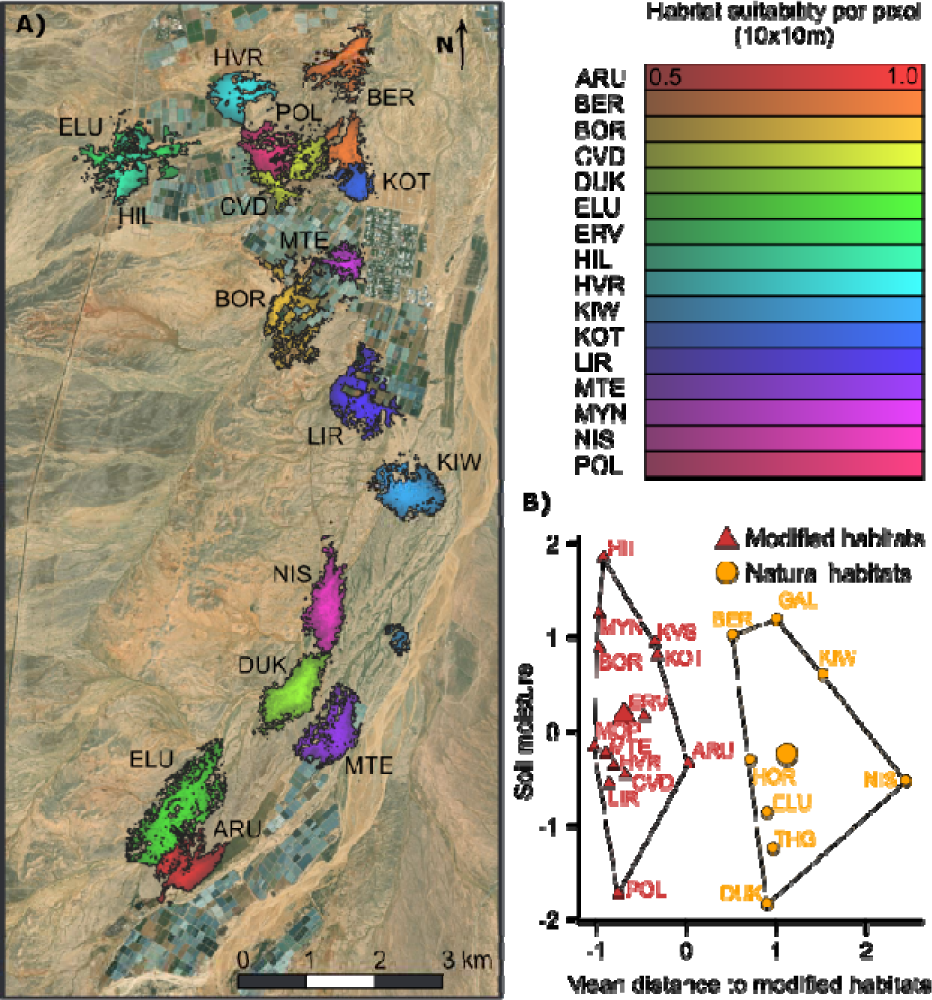
**A) Distribution models projected on the study area for April 2021.** Each polygon represents the distribution model for each of the study groups. The color gradient represents the values of suitability from 0.5 (darker colors, less suitable) to 1 (lighter colors, more suitable). **B) Classification of the study groups in clusters based on average values of distance to modified habitats and humidity of territories.** Groups from natural habitats are represented by orange circles and groups from modified habitats by red triangles. The center of the cluster is represented by a larger symbol.

#### Environmental clustering and habitat classification

We divided the 21 study groups by k-means clustering into two habitats: Modified and Natural (compactness = 81 %, average silhouette width = 0.73; Fig. 2B). Modified territories comprised all groups with territories closer to anthropogenically modified habitats (N = 13; average distance ± SE = 145.67 ± 36.18 m) and natural territories comprised all groups with territories that are remote from modified habitats (N = 9; 886.67 ± 87.92 m). Average values of soil moisture were similar within both categories (modified: −20.50 ± 0.31, natural: −20.03 ± 0.46).

### Fitness

#### Breeding events, breeding success and nesting success

Our analyses revealed a strong negative effect of natural habitats on the number of breeding events per season (estimate ± SE, −0.466 ± 0.155; p = 0.002; Fig. 3A). Groups from natural habitats had fewer breeding events than groups from modified habitats (average ± SE; natural: 1.58 ± 0.18, modified: 2.47 ± 0.14). Models that included total rainfall were the best, based on corrected Akaike Information Criterion (AICc) values. Total rainfall was positively related to the number of breeding events in both habitats (estimate ± SE, 0.010 ± 0.003; p = 0.004, Supplementary Information Fig. 1A). We found moderate evidence that habitat affected the number of fledglings raised per breeding season, with a negative effect of natural habitats (estimate ± SE; −0.670 ± 0.273; p = 0.014; Fig. 3A). On average, groups from modified habitats raised more fledglings than groups from natural habitats (average ± SE; natural: 3.00 ± 0.55, modified: 4.55 ± 0.42). We also found a very strong evidence that total rainfall together with the number of adults and subadults (individuals which were born in previous breeding season and act as helpers but are not older than one year) in a group had a positive effect on the number of fledglings raised (estimate ± SE; total rainfall: 0.022 ± 0.005; p < 0.001; adults and subadults: 0.097 ± 0.030; p = 0.001; Supplementary Information Figs. 1B,1C). No relationship was found between nesting success and habitat (p = 0.225, Fig. 3A); however, we found moderate and strong evidence that total rainfall and adults and subadults and total rainfall had a positive effect on nesting success (estimate ± SE; total rainfall: 0.017 ± 0.005; p = 0.033; adults and subadults: 0.133 ± 0.055; p = 0.009; Supplementary Information Figs. 1D,1E).

**Figure 3.**
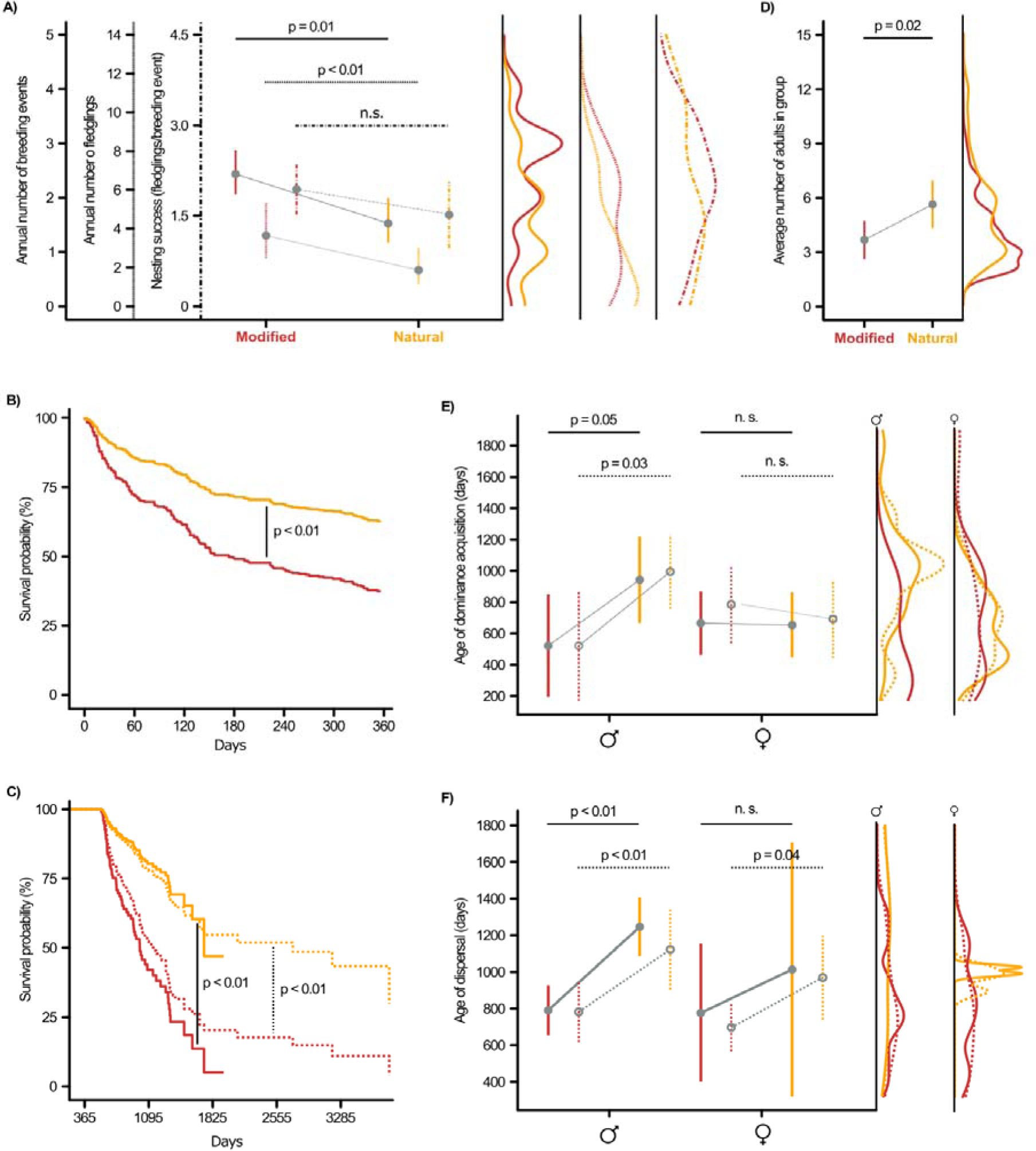
**A) Effect of habitat on the number of breeding events performed by groups per year (full line), the number of fledglings raised per year (pointed line), and nesting success (fledglings to breeding events ratio; pointed-dashed line); B) Cox hazard model for juvenile survival. C) Cox hazard model for adult survival full line represents the model prediction with the restricted dataset and pointed line represents the model prediction with wide dataset; D) Effect of habitat on the average number of adults in the group per month; E) Effect of habitat on the age of dominance acquisition in males and females in the restricted dataset (full line) and in the wide dataset (dashed line); F) Effect of habitat on the age of dispersal in males and females in the restricted dataset (full line) and in the wide dataset (dashed line).** The linear graphs on the right represent the frequency of observations on the raw data for each sex and dataset. In all the graphs, natural habitats are represented in orange and modified habitats in red. In the significative effects the p-values are shown. Non-significant effects are marked as ‘n.s.’

#### Juvenile survival

In total, 136 out of 373 juveniles survived to adulthood (36.46 %). Of them, 82 out of 275 were from modified habitats (29.81 %), and 54 out of 98 were from natural habitats (55.10 %). We found very strong evidence that juveniles in natural habitats were more likely to survive to their first year of life than those in modified habitats (estimate ± SE, −0.746 ± 0.195, p <0.0001; Fig. 3B). Survival probability to adulthood was also higher in natural (predicted, confidence interval: 0.596, 0.505 - 0.702) than in modified habitats (0.337, 0.285 - 0.398) also when compared with fledging to independence (natural: 0.84, 0.789 - 0.894; modified: 0.693, 0.644 - 0.747). The number of adults present in the group during the first 60 days of life did not affect juvenile survival (0.082 ± 0.04 p = 0.074), nor did the fledglings-adults ratio at the time of hatching (0.304 ± 0.165p = 0.065), brood size (−0.149 ± 0.089, p = 0.095), the number of former breeding attempt (−0.118 ± 0.099, p = 0.233), or the average NDVI (3.274 ± 2.156, p = 0.128).

#### Adult survival

We used two datasets to assess adult survival: the restricted dataset, which included only the adults born during the study dates, and the wide dataset, which included all the adults present in the study groups during the study period regardless of age. We found very strong evidence that adult babblers in natural habitats lived longer than those in modified habitats in the restricted dataset (−1.375 ± 0.335, p < 0.0001; Fig. 3C) and in the wider dataset (−0.972 ± 0.378, p = 0.0001; Fig. 3C). In the restricted dataset, the survival rate of adults to the second year of life in natural habitats was higher (predicted, confidence interval: 0.880, 0.811 - 0.955) than in modified habitats (0.605, 0.507 - 0.720), as well as for the third year of life (natural: 0.772, 0.658 - 0.906; modified: 0.360, 0.256 - 0.506) and the fourth year (natural: 0.603, 0.430 - 0.846; modified: 0.135, 0.052 - 0.349). Similar results were found for the wider dataset: second year (natural: 0.882, 0.826 - 0.941; modified: 0.717, 0.641 - 0.802), third year (natural: 0.794, 0.711 - 0.887; modified: 0.544, 0.456 - 0.649), and fourth year (natural: 0.749, 0.653 - 0.858; modified: 0.466, 0.374 - 0.580). In the restricted dataset, we found weak evidence of the effect of sex in the survival probabilities for adults (0.490 ± 0.271, p = 0.070) (Supplementary Information Table 3). In the wider dataset, sex did not affect survival (0.124 ± 1.132, p = 0.578).

#### Group size

Our analysis revealed a moderate positive association between natural habitats and group size (estimate ± SE, 1.962 ± 0.811; p = 0.026; Fig. 3D). Groups from modified habitats were smaller than groups from natural habitats (average number of individuals ± SE, modified: 3.90 ± 0.06; natural: 5.19 ± 0.12).

### Life history traits

#### Dominance acquisition

In the restricted dataset, we found that males living in natural habitats attained first dominance later in life than those in modified habitats (estimate ± SE, 456.07 ± 98.82; p < 0.001; Fig. 3E). Males from natural habitats acquired dominance at an average age (±SE) of 1171.33 (± 175.10) days, while males from modified habitats acquired dominance at an average age (±SE) of 807.53 (± 65.46) days. We also found that in both habitats, the number of adult males in the group delayed the age of dominance acquisition (estimate ± SE, 261.40 ± 41.32; p < 0.0001). In the wider dataset, we found similar results for the effect of modified habitats on the age of dominance acquisition (estimate ± SE, 340.10 ± 133.70; p = 0.023; Fig. 3E). Males from natural habitats acquired dominance at an average age (±SE) of 1113.30 (± 130.11) days, while males from modified habitats acquired dominance at an average age (±SE) of 783.05 (± 53.64) days.

There was no evidence of the effect of habitat (p = 0.490) or the number of adult females in the group (p = 0.679; Fig. 3E) on the female age when first attaining dominance in the restricted dataset. In the wider dataset, we found moderate evidence that females from modified habitats attained dominance at a younger age than those from natural areas (estimate ± SE, 272.73 ± 122.10; p = 0.047; Fig. 3E). Females from natural habitats acquired dominance at an average age (±SE) of 976 (± 28.04 days), while females from modified habitats acquired dominance at an average age (±SE) of 713.36 (± 66.67) days.

#### Delayed dispersal

The analysis of the dataset restricted to small coalitions and non-forced dispersal showed weak evidence that males in natural habitats dispersed later than males from modified habitats (estimate ± SE, 421.9 ± 191.3; p = 0.051; Fig. 3F). Males in natural habitats dispersed for the first time at an average age (±SE) of 943.14 (± 122.34) days, while males in modified habitats dispersed at an average age (±SE) of 521.20 (± 148.04) days. The full dataset that included all of the individuals that dispersed during the study period showed that males in natural habitats dispersed later than males from modified habitats (estimate ± SE, 473.4 ± 197.9; p = 0.031; Fig. 3F). Males in natural habitats also dispersed for the first time later, at an average age (±SE) of 994.54 (± 114.68) days, while males in modified habitats dispersed at an average age (±SE) of 521.20 (± 148.04) days. We found no effect of habitat on the age of first dispersal in females in the restricted dataset (p= 0.922) nor in the wider dataset (p = 0.590, Fig. 3F).

## Discussion

Our results indicate a consistent, strong, negative influence of modified habitats on the fitness, life history, demography, and behavior of a desert living, cooperatively breeding system. Groups in modified habitats had more breeding events per season but lower juvenile and adult survival, smaller groups, and lower group stability. This may be in part due to earlier male dispersal and dominance attainment in modified habitats, while they are less experienced. Thus, habitat changes affect the life history, fitness, cooperation, and group decision-making processes. Our findings suggest that modified habitats may serve as an ecological trap for native species, especially in arid environments where resources are scarce.

Smaller groups producing smaller dispersing cohorts may have lower survival rates or lower dispersal success, driving populations of cooperative breeders to extinction^33^. It is also possible that human interferences, such as habitat degradation and persecution, reduce the number of potential natural patches that are suitable for colonization as well as the maximum group size for each patch, eventually leading to a demographic Allee effect^33,37^. Empirical and theoretical studies support this hypothesis^42,43^, including in the studied population of Arabian babblers^37^, which is confirmed here, with strong evidence of the effect of habitat modification on cooperative breeder fitness.

The agricultural areas in the Arava Valley introduced an abundance of resources into a naturally hyper-arid environment and increased the availability of water and food, starting in the late 1960s. Arabian babbler groups responded to the modification process by moving towards the modified areas and increasing their breeding events in them, even during suboptimal periods such as the dry, hot months of summer, where daytime temperatures reach ∼ 45°C. Environmental changes that increase the availability of resources over time are known to extend the breeding season and the number of breeding attempts per season in various bird species^10,31,44^. The increase in the number of breeding attempts, with similar fledging success rates between groups in natural vs. modified habitats, causes a higher fledgling productivity per year rate in modified habitats. Despite this higher productivity, the recruitment rates were lower for the latter due to lower juvenile survival during their first year of life. In Arabian babblers, the highest mortality rates occur between fledging and independence^39^. While the juvenile survival rates in natural habitats were similar to rates of other cooperatively breeding species^45–47^, it was lower in modified habitats. We suggest two possible explanations for this: 1. Juveniles are less adapted to the modified habitats and require more time to learn the necessary foraging skills^48,49^; Juveniles are also more explorative and risk-prone than adults^50^, putting them at increased predation, poisoning, or harm risks in an unfamiliar environment. 2. The extent of adult care in cooperative breeders is crucial for juvenile survival^39,51,52^, but in modified habitats, adult care may be reduced due to smaller groups and earlier dispersal (findings from this study).

A direct consequence of the negative effect of modified habitats on adult survival is the increased availability of dominant positions in groups, which could facilitate an earlier dispersal based on ecological constraints^29,53^ or benefits of philopatry hypotheses^53,54^. Dispersal is a costly behavior for cooperative breeders^40,53^, and the presence of smaller groups that are more likely to accept new members until the optimal group size is reached^26^, may incentivize dispersal in modified habitats. Surprisingly, although these cues would be expected to favor dispersal in both sexes, our results showed it only in the non-dispersing sex, males, and not in females. Given that we did not find sex differences in survival, the simplest explanation is the documented sex differences in dispersal strategies in the Arabian babbler. In this species, males tend to remain in their natal group^55^. The results of the current study suggest that dispersal-related decisions are affected by the increased breeding position availability due to increased adult mortality rates in modified habitats. Similar results were found in pied babblers (*Turdoides bicolor*), where breeding vacancy availabilities increased the dispersal probabilities of males but not of females^53^, although this species dispersal is not sex-biased^56^. In white-winged choughs (*Corcorax melanoramphos*), dispersal was sex-biased towards females only in drought years, and increased mortality promoted female dispersal, but not males’^34^.

Reduced survival in modified habitats may also be related to faster turnover of dominant positions and consequently the younger age for attaining dominance. Younger and less experienced dominant individuals negatively affect the breeding success of groups in other cooperative breeding species^25,57^. Therefore, our findings suggest that the effects of habitat on sociality might enhance its effect on fitness. On the other hand, negative effects on fitness can also have consequences on sociality; for example, smaller groups and shorter social queues to attain dominance. In this sense, we expected similar results for both sexes in age of dominance acquisition, but we found that only male status was related to habitat. Nevertheless, when we studied the wide dataset, we found evidence of a similar trend in females. Hence, the small female sample size in the restricted dataset (n = 7) may cause an underestimation of the effect of habitat on females.

This study revealed that modified landscapes in arid ecosystems entail a source-sink scenario for a cooperative breeder and desert specialist, due to the low rates of recruitment in modified habitats together with high adult mortality rates. Moreover, we consider that these agricultural landscapes imply also a potential ecological trap for Arabian babblers. The perception of modified habitats as high-quality habitats may have led Arabian babbler groups to attempt to breed more and attain higher productivity. Nevertheless, this perception was found to be misleading, and the actual breeding performance, lower survival rates, and recruitment in these new habitats proved otherwise. To label modified habitats an ecological trap, the increased attractiveness of these newly created habitats over natural habitats must be proven. Indeed, our preliminary results suggest higher group densities in modified than in natural habitats and an ongoing movement of groups from natural to modified habitats (A. Alamán, unpublished data). To explore these spatial and temporal changes in the delimitation of anthropogenically-modified habitats, remote sensing has been found as a useful tool, providing a reliable and repeatable source of information for habitat characterization^58^. The use of species distribution models to infer Arabian babbler group territories is novel, and helped us understand how anthropogenic habitat transformation affected population distribution. In the future it can be used to model other types of processes, for example climate change.

In conclusion, in this study we showed how human habitat modification are negatively related to cooperative breeders’ fitness and life history traits. The outcome includes smaller groups^33^, longer breeding seasons^31^, and lower juvenile survival^32,34^, which are potentially detrimental. Habitat effects go beyond fitness and breeding performance, including deeper aspects of sociality and individual decisions, which are affected by habitat modification. Group social and structural changes could increase the fitness consequences of habitat modification and lead the population to collapse. Amid the global change scenario that is currently unfolding, which includes rapid and extreme ecosystem changes, species’ degree of sociality, life history, and relationship with habitat should be considered to develop effective conservation plans.

## Methods

### Study area and field data collection

Data was collected between 2016 and 2021 from the population of Arabian babbler in the Sheizaf Nature Reserve (30° 43 ‘N, 35° 15 ‘E), an extremely arid desert in the Arava Valley (Israel; Fig. 4A). The reserve is a 52.5 km^2^, divided into the north (6.5 km^2^) and the south (46 km^2^) protected areas. Numerous dry riverbeds (i.e., wadis) cross the reserve and are where the natural vegetation is concentrated, mainly Umbrella thorn acacias and shrubs^39^. Four agricultural settlements are at the reserve’s borders: Idan in the northeast, Ein-Hazeva in the northwest, Hazeva between the north and south protected areas, and Ein Yahav in the south. Near these settlements, the landscape has been transformed into farmland, where two kinds of cultivation techniques are used: open-air crops (mainly date and mango plantations) and greenhouses^15^.

**Figure 4.**
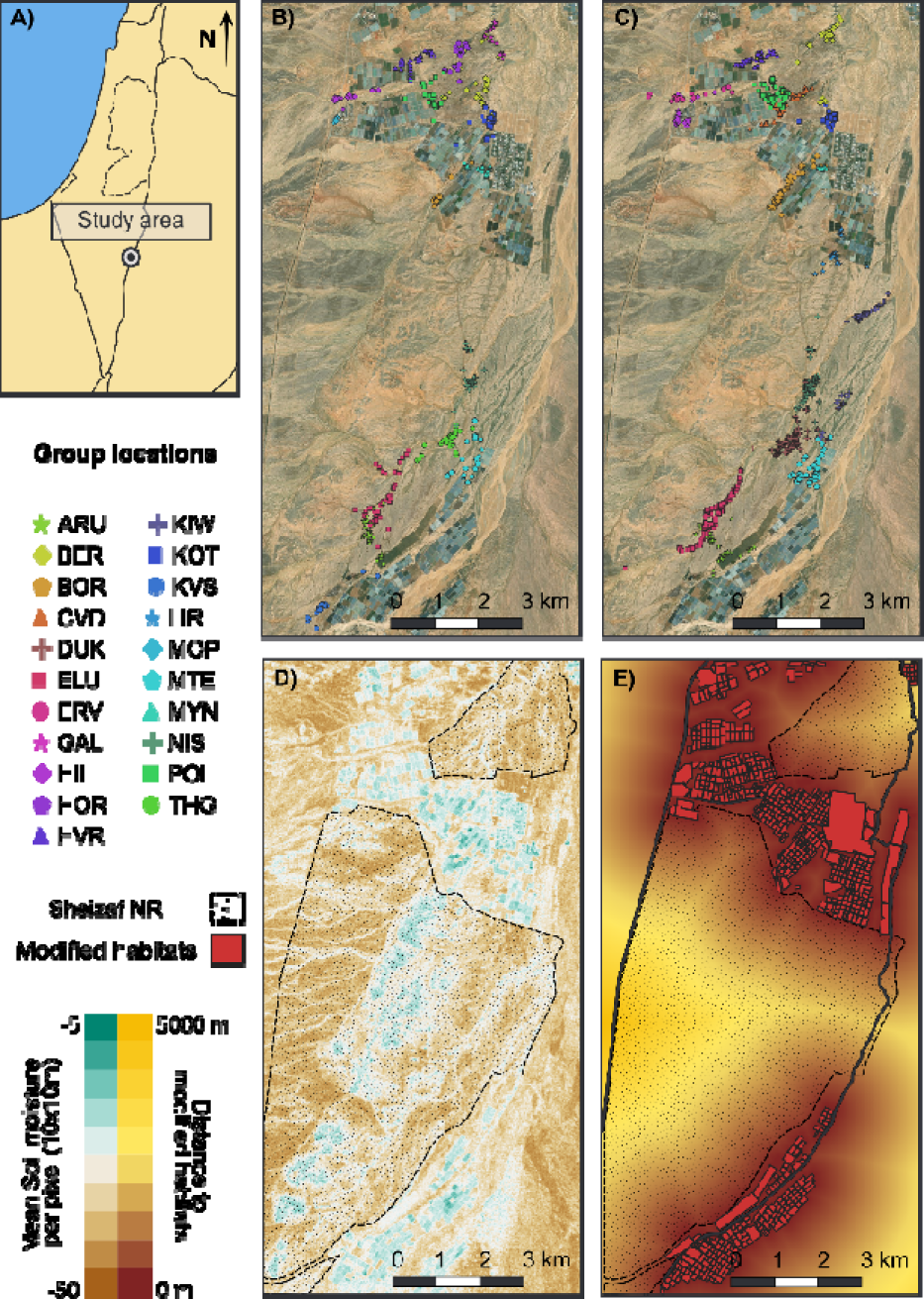
**A) Location of the study area within Israel; B, C) Study group locations during 2020 and 2021 breeding season (respectively)**. Symbols and colors represent the GPS location for each of the study groups; **D) Map of soil moisture per 10 x 10 m pixel in March 2021 extracted from Copernicus Sentinel 1 (S1) imagery (https://developers.google.com/earth-engine/datasets/catalog/COPERNICUS_S1_GRD)**. Blue corresponds to higher and brown to lower soil moisture values; **E) Heat map of the Euclidean distances to modified habitats crops, greenhouses, and settlements) derived from maximum NDVI extracted from Sentinel 2 (S2) imagery (https://developers.google.com/earth-engine/datasets/catalog/COPERNICUS_S2_SR_HARMONIZED)**. Yellow-red gradient indicates distance to modified habitats (yellow more distant, red more proximal).

The study population of Arabian babblers is in the natural protected areas of the reserve as well as in the modified habitats (e.g., farmland and settlements). The research was established by Amotz & Avishag Zahavi in 1971 and has been studied continuously to the present^38^. All individuals in the study groups are ringed with unique combinations of color rings that allow individual recognition, and birds are habituated to the presence of observers. Sex is determined visually once individuals are mature (> one year old), by the color of iris (yellow in males, dark brown in females). The study population is monitored daily. The location of the group and the number of adult, subadult (individuals that were born in the previous breeding season and acted as helpers but were not older than one year) and juvenile individuals are collected using the application Anecdata.org^59^. Any change in the status of an individual (e.g., change in hierarchy, dispersal events, disappearance, or death) is entered into a database. Daily monitoring during the breeding season includes a search for new nests and monitoring active nests. The number of eggs, nestlings, and any change in nest condition are noted for each nest and entered into the database. If direct nest observations are not possible, egg-laying date is inferred by the incubation activity of the dominant pair, and egg-hatching dates are estimated by the beginning of food provisioning by the group to the nestlings. Nestlings are ringed nine to ten days after hatching^60,61^. Further information about the research population, habituation, and ringing protocols can be found in ref.^62^. Permits for this study were granted by the Israel National Parks Authority (permit numbers: 2022/43151, 2020/42538, 2018/41848, 2016/41453).

### Habitat characterization

We considered 4665 locations collected during the 2020 (1499 locations, Fig. 4B) and 2021 (3166 locations, Fig. 4C) breeding seasons for 21 different groups. We only included study groups those with at least 30 locations collected during the breeding season. In order to identify habitat preferences of different groups of babblers towards anthropogenically modified or natural areas, two steps were followed: (i) Species distribution modelling and (ii) Environmental clustering and Habitat classification.

#### Species Distribution Models

A series of Species Distribution Models (SDM) was constructed to identify the spatial distribution of the different groups, considering a series of environmental variables derived from freely available RS products and images from the Copernicus Earth Observation program of the EU (https://land.copernicus.eu/, accessed on 15/04/2022), both optical and radar. Along with these environmental variables, we considered the collected groups’ locations for 2020 and 2021. On the one hand, Copernicus Sentinel 1 (S1) imagery (https://developers.google.com/earth-engine/datasets/catalog/COPERNICUS_S1_GRD) was used to retrieve information regarding soil moisture in the study area^63^. The Synthetic Aperture Radar (SAR) backscatter VH coefficient of this Remote Sensing (RS) imagery was used as a proxy for soil moisture ^64^ (Supplementary Information Fig. 4D). Sentinel 2 (S2) imagery (https://developers.google.com/earth-engine/datasets/catalog/COPERNICUS_S2_SR_HARMONIZED) was used to assess vegetation density using Normalized Vegetation Index (NDVI) using bands 4 (red) and 8 (near infrared)^65^. Google Earth Engine (GEE)^66^ was used to calculate monthly average values of SAR VH backscatter and NDVI for months corresponding to the Arabian babbler breeding season (March-August) between 2017 – 2021. We selected the breeding season for habitat characterization because during this period group territories are most stable^38^.

Cultivated areas and anthropogenic vegetated areas were identified by studying the maximum NDVI. We considered NDVI values higher than 0.7 as densely vegetated areas^67^ to identify cultivated and anthropogenic patches (hereafter modified habitats) by visually interpreting their spatial structure (e.g. regular patterns of NDVI were associated with crops). Euclidean distance to cultivated areas was calculated using QGIS v3.22 Białowieża^68^ (Fig. 4E). All habitat data was projected to WGS 84/UTM zone 36N Coordinate Reference System, with a final spatial resolution of 10 m x 10 m.

#### Environmental clustering and habitat classification

Considering the identified spatial distribution of the species, the environmental preferences of each group were identified, depicted by environmental characteristics such as (i) Euclidean distance to previously identified modified habitats and (ii) soil moisture. Those values were calculated using weighted average values based on groups’ suitability, so that when calculating this average, the weight of environmental variables correspond to the number of pixels and suitability. Once environmental preferences were assessed, we used the average weighted values of soil moisture and distance to modified habitats of the territories to perform a k-means analysis^69^ and classify the groups as modified or natural based on the similarity of those preferences.

### Fitness

#### Breeding events, breeding success, and nesting success

We counted the total number of breeding events per season. A breeding event was established once an egg was laid in the nest. To assess breeding success, we counted the total number of individuals who fledged during the season per group. Nesting success was calculated as the number of fledglings divided by the total breeding events in the breeding season.

As rainfall has been found to be an important factor for breeding success and survival in other cooperative breeder species in extremely arid habitats^70^, we calculated three different measurements of rainfall as environmental factors for each of the study years. Total rainfall from October of the previous year to April of the study year, winter rainfall as total rainfall from October of the previous year to February of the study year, and breeding rainfall as total rainfall from March to August of the study year. All rain variables were calculated in total millimeters of rainfall. Climatic data were extracted from Israel meteorological services (https://ims.gov.il/en/data_gov)

#### Juvenile survival

We included in this analysis only ringed individuals who hatched in the study site during the study period and their exact age was known. We calculated the age of the individuals when they died, dispersed or the study period ended (31/08/2022). An individual was considered dead once it was not observed in the study area. Since long distance dispersal cannot be distinguished from death, the surrounding of the study area were also monitored to minimize mortality overestimation^53^. We also calculated the average number of adults and subadults during the first 60 days after fledgling^39^, the fledglings-adult ratio at the time of hatching, brood size (number of individuals hatched at the nest), number of breeding events in the breeding season, and average NDVI of the territory at the month of hatching. Sex is not visually distinguished in individuals younger than one year. In total, 373 (275 from modified habitats and 98 from natural) juveniles were included in the analysis.

#### Adult survival

The restricted dataset (N = 135 adults; 68 males, 67 females) included ringed individuals that fledged and reached adulthood (i.e., one year old) during the current study. We only included individuals who remained in the same group from birth to their first dispersal or last sighting to keep the effect of habitat to those that are affected during their entire lifespan. Like in juveniles, an individual was considered dead once it was not observed in the study area. To avoid overestimation of mortality, the individuals that disappeared in the last two months of the study (n = 10) were considered as unknown status.

The full dataset (N = 182; 99 males, 83 females) included all adults present in the study groups during the study period, no matter when they were born. Dataset was still restricted to individuals who remained in their natal groups until the study period to keep habitat as a constant effect.

#### Group size

The group size was calculated as the average monthly number of adults in the group. Dates of entry and departure to and from a group were based on daily observations. An individual was considered a part of a group once it was observed roosting overnight with that group for at least three consecutive nights.

### Life history traits

#### Dominance acquisition

We used GLM to analyze the effect of habitat on individual age (in days) of first attaining a dominant rank. We only included individuals who hatched in the study site and attained dominance during the study period (N = 28 individuals; 19 males, 7 females). We determined the first age as dominant once the individual replaced the previous dominant (either through chasing the dominant or through replacing a dominant that disappeared). We created one additional datasets (N = 44; 29 males, 15 females), which included all the dominant individuals who were present during the study period, regardless of when they attained dominance.

#### Delayed dispersal

A dispersal event was determined once an individual left their natal group and was observed joining a new group or acting as a floater. We included all the individuals who hatched and dispersed for the first time during the study period (N = 37; 12 males, 25 females). We excluded individuals who formed coalitions of three or more and dispersed together and individuals who were actively chased away from the group. We repeated the analysis with a wider dataset where these individuals were included (N = 57; 16 males, 41 females).

### Statistical analysis

All statistical analyses were performed within r v4.0.3^71^. We used Generalized Additive Models (GAM)^72^ with Biomod 2 v3.3.7 r package for Species Distribution Models (SDM)^73^. K-means analysis was performed with the package ‘stats’ v4.0.3^71^. We used packages ‘lme4’ v1.1.27^74^ for Generalized Linear Mixed Models (GLMMs)^75^ analysis and ‘survival’ v3.3^76^ for multivariate survival analysis (Cox proportional-hazards models^77^). We used package ‘effects’ v4.2^78^ for display the variables’ effect in our models and plotted it with ‘graphics’ v4.0.3^71^. For drawing survival curves, we used package ‘survminer’ v0.4.9^79^. Maps were represented with QGIS v3.22. All other figures were drawn with ‘ggplot2’ v3.3.5^80^.

GAM was selected to construct our distribution model, as it has been found to present particularly robust performance in SDM^81,82^. A five-fold cross-validation technique with 25% test samples was used^83^. Model performance evaluation criteria were based on: (i) Area Under the Curve (AUC)^83^ and (ii) Model specificity and sensitivity^84^. The final projection of the model to the study area provided a per-pixel group suitability, getting values from 0 (no suitability) to 1 (perfect suitability), allowing identifying the spatial distribution of each group.

For breeding events analysis, we constructed three different GLMMs, each with one of the rainfall variables and habitat as explanatory variables. Group identity and year were included as random factors. For breeding attempts, a Poisson distribution of errors was used. For breeding success, we also constructed three different GLMMs with the different rainfall variables and habitat and number of adults and subadults during the breeding season as explanatory variables. The total number of fledglings was used as a response variable. Group and year were included as random factors. We set a Poisson distribution of errors for this analysis. Finally, we performed another set of three GLMMs with the different rainfall variables, with habitat and number of adults and subadults and ratio between fledglings and total number of breeding events to assess nesting success between habitats. Group and year were included as random factors. In all the analyses, model fit was assessed by Akaike’s Information Criterion for small sample sizes (AICc)^85^, and we selected the best model based on these values (Supplementary Information, Table 1).

For juvenile and adult survival, we used Cox proportional-hazards models to explore the payoffs from different habitat choices at a local scale. Though observation-based survival analysis on wild populations might overestimate mortality rates when considering permanent long-distance dispersal as deaths, it is still an accurate tool to estimate payoffs from different habitat choices^53^. For each individual, we included the number of days that it was observed and if an event (i.e., death) occurred during this period or not. If we stopped following an individual before the period of study was finished (if the individual dispersed or had an unknown status), it was included as censored data, so that the model considers these individuals for calculating the survival likelihood. For juvenile survival analysis we included as explanatory variables: habitat, the average number of adults and subadults in the group during the first 60 days after fledgling, the fledglings-adults ratio, brood size, number of breeding events, and average NDVI. In adult survival analysis, we also included habitat and sex as explanatory variables.

We used GLMMs to assess how habitat affected group size. We included habitat as the explanatory variable with group identity and month nested within year as random factors.

For the analyses of effect of habitat on age of dominance acquisition and first dispersal we built separated GLMMs for each sex, with habitat as the explanatory variable, and age at first attaining dominance or dispersal as the response variable. For dominance analysis, the number of same-sex adults was also included as an explanatory variable in the restricted datasets. Group identity was included as a random factor.

We used the *cox.zph* function of the package *survminer* to test if our models fit the proportional hazards assumption. The function correlates the scaled Schoenfeld residuals of each variable with the time. Non-significative results of this test refuted the assumption of a relationship between time and the variables (Supplementary Information Table 2).

## Supplementary information

**Figure 1.**
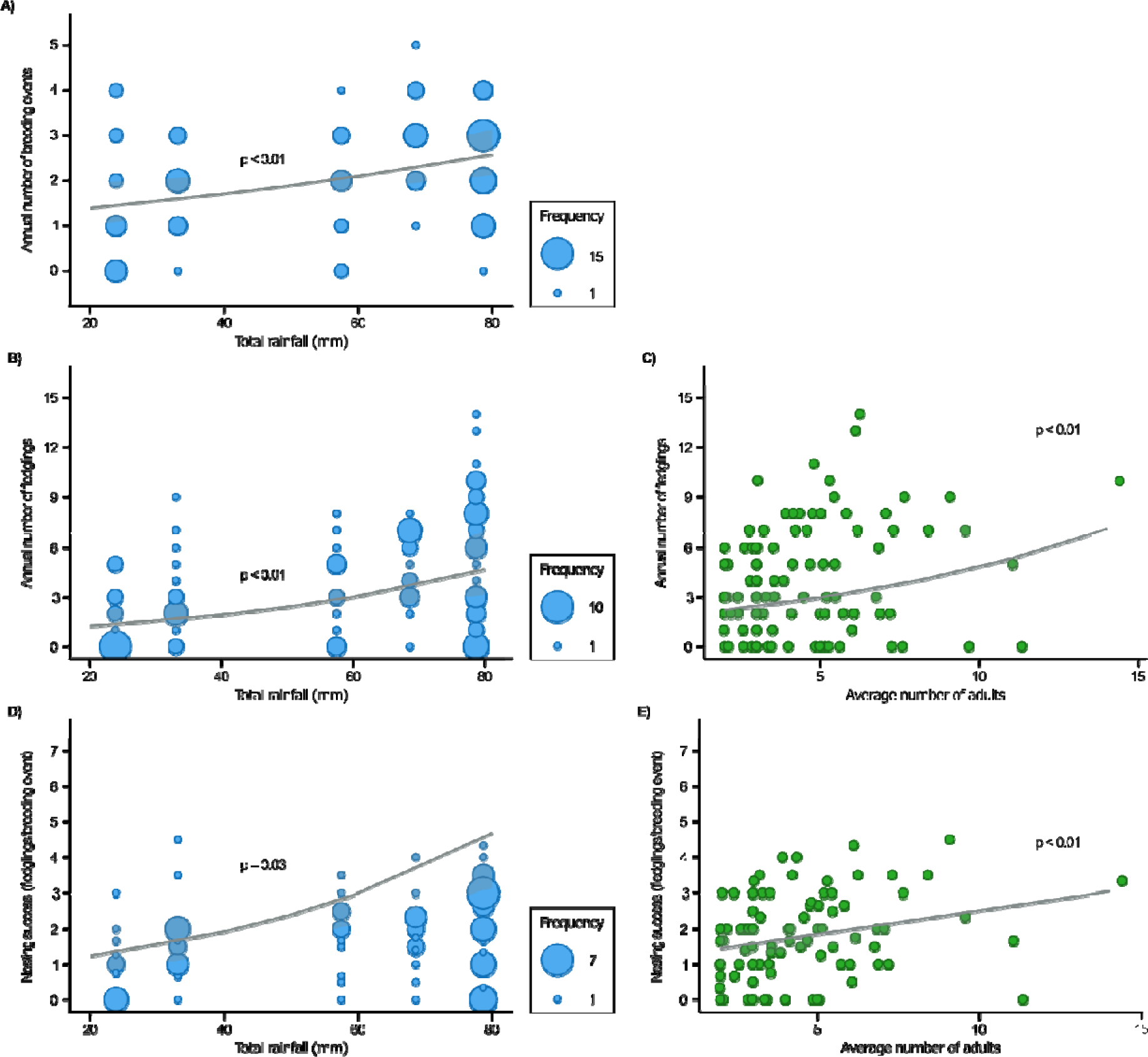
**A) Effect of total annual rainfall (mm) on the number of breeding events per group per breeding season. B). Effect of total annual rainfall (mm) on the number of fledglings raised per group per breeding season. C) Effect of average number of adults and subadults on the number of fledglings raised per group per breeding season. C) Effect of total annual rainfall (mm) on the ratio between the number of fledglings per total number of breeding attempts per group per breeding season; b) Effect of average number of adults on the ratio between the number of fledglings per total number of breeding attempts per group per breeding season**. Shaded areas show the 95% confidence interval. Points represent the raw data during the study period.

**Table 1.** Summary of the generalized linear mixed models generated for analysis of number of attempts, breeding success and nesting success. Asterisks indicate the significative explanatory variables. AICc shows the value of Akaike’s Information Criterion for the full model. AICc 2 shows the value of Akaike’s Information Criterion for the simplified model with only significative variables. Bold models indicate the selected models in our study.

| Response variable | Explanatory variables | Random factors | AICc | AICc 2 |
| --- | --- | --- | --- | --- |
| **Attempts** | **Habitat\***<br>**Number of adults and subadults**<br>**Total rainfall\*** | **Group**<br>**Year** | 310.016 | **309.195** |
| Attempts | Habitat*<br>Number of adults and subadults<br>Winter rainfall | Group<br>Year | 314.268 | 312.734 |
| Attempts | Habitat<br>Number of adults and subadults<br>Breeding rainfall | Group<br>Year | 313.867 | 312.734 |
| **Number of fledglings** | **Habitat\***<br>**Number of adults and subadults\***<br>**Total rainfall\*** | **Group**<br>**Year** | **435.669** |  |
| Number of fledglings | Habitat*<br>Number of adults and subadults*<br>Winter rainfall | Group<br>Year | 442.78 | 441.476 |
| Number of fledglings | Habitat*<br>Number of adults and subadults*<br>Breeding rainfall* | Group<br>Year | 440.796 |  |
| **Breeding success** | **Habitat**<br>**Number of adults and subadults\***<br>**Total rainfall\*** | **Group**<br>**Year** | 285.071 | **283.905** |
| Breeding success | Habitat<br>Number of adults and subadults*<br>Winter rainfall | Group<br>Year | 286.791 | 279.294 |
| Breeding success | Habitat<br>Number of adults and subadults*<br>Breeding rainfall | Group<br>Year | 289.11 | 279.294 |

**Table 2.**
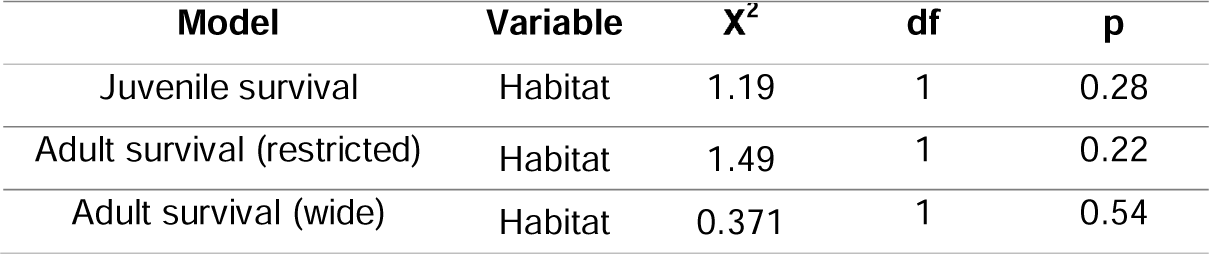
Summary of the test of proportional hazard assumption in the survival models.

| Model | Variable | X<sup>2</sup> | df | p |
| --- | --- | --- | --- | --- |
| Juvenile survival | Habitat | 1.19 | 1 | 0.28 |
| Adult survival (restricted) | Habitat | 1.49 | 1 | 0.22 |
| Adult survival (wide) | Habitat | 0.371 | 1 | 0.54 |

**Table 3.**
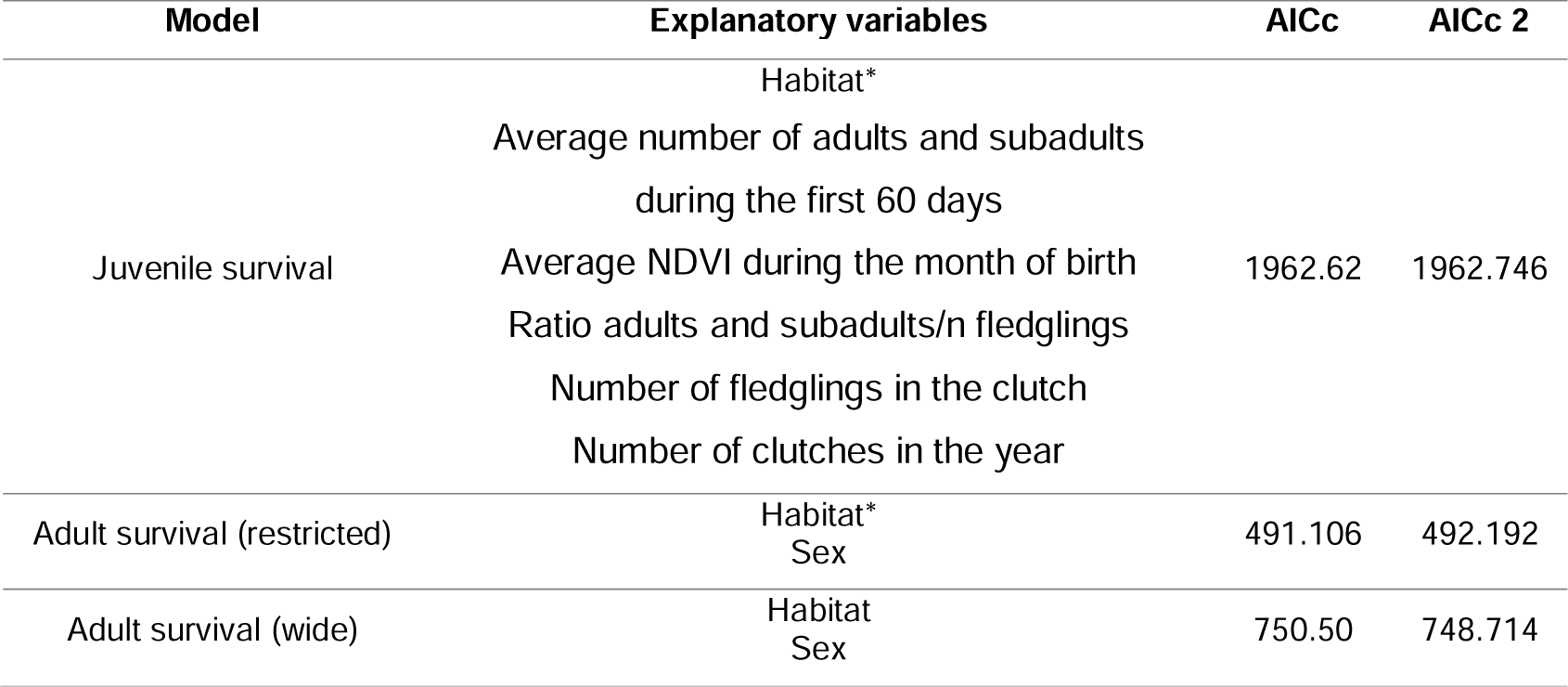
Summary of the Cox proportional hazard models generated for analysis of juvenile and adult survival. Asterisks indicate the significative explanatory variables. AICc shows the value of Akaike’s Information Criterion for the full model. AICc 2 shows the value of Akaike’s Information Criterion for the simplified model with only significative variables.

